# eDNA on the go: a direct comparison of fixed and vehicle mounted airborne eDNA sampling methods for terrestrial vertebrate species detection at large spatial scales

**DOI:** 10.1101/2025.09.02.673667

**Authors:** Francesca Martino, Christine Cooper, Josh Newton, Philip Bateman, Josh Kestel, Mieke van der Heyde, Morten Allentoft, Mahsa Mousavi-Derazmahalleh, Angus Lawrie, Austin Guthrie, Paul Nevill

## Abstract

Air is receiving increasing recognition as a biologically rich source of taxonomically diverse environmental DNA (eDNA). Multiple proof-of-concept studies have explored air as a medium for the detection of single terrestrial species, and even entire terrestrial communities, including mammals, non-anemophilous plants and insects. Airborne eDNA has been sampled using various stationary devices but if we can access it using mobile collection methods, we can rapidly assess biodiversity at large spatial scales. We therefore compared passive eDNA filters deployed for short (30 mins) and long periods (24 hrs) with filters attached to cars that sampled transects through natural and human-modified environments. Our metabarcoding procedure detected 49 vertebrate taxa (33 birds, 15 mammals and 1 amphibian) from all three eDNA air sampling approaches combined, with 29 of these taxa detected in samples collected using the mobile method. The total number of taxa, or proportions of unique taxa detected, did not differ by sampling method, land use or day. Community composition was not significantly impacted by sampling method or sampling day but differed significantly between land uses. We propose that our car-mounted air eDNA sampler could be a game changer for large scale, rapid biomonitoring efforts.

## INTRODUCTION

Biodiversity is increasingly threatened by anthropogenic impacts including habitat clearing and fragmentation (Brodie et al., 2021; Engert et al., 2023), pollution, climate change (Barnosky et al., 2012; Butchart et al., 2010; Li et al., 2016) and the introduction of invasive or domestic species (Butchart et al., 2010; Cardinale et al., 2012). These threats not only negatively influence individual species, but also disrupt complex ecological networks, leading to cascading effects throughout ecosystems (Ceballos et al., 2017). Vertebrate species in particular are facing significant declines (Ceballos et al., 2017; Li et al., 2016), but maintaining vertebrate diversity is essential for maintenance of ecosystem services such as pollination, pest control, seed dispersal, and the balance of food webs (Ceballos et al., 2017). Vertebrate species are also fundamental for maintaining the cultural, aesthetic and ethical values that human society places on wildlife and nature (Schultz et al., 2005).

Biodiversity monitoring (hereafter referred to as biomonitoring) is the crucial first step in conservation and restoration efforts (Campbell et al., 2002; Niemelä, 2000). It provides the essential data needed to assess populations, ecosystem health and identify areas most in need of intervention (Capdevila et al., 2022). For animals, this information is typically acquired by physically identifying species in their natural habitats (Baird & Hajibabaei, 2012; Ruppert et al., 2019). While effective and frequently applied, conventional field survey approaches such as camera traps, transect surveys, quadrat sampling, live-trapping and mist-netting require significant time and resources to implement (Baird & Hajibabaei, 2012), potentially negatively affect wildlife (Thomsen & Willerslev, 2015) and and can involve considerable bureaucracy associated with occupational health and safety (OHS) and animal ethics requirements. They also require taxonomic expertise to identify species (Deiner et al., 2017) and can underestimate the presence of cryptic species (Hending, 2024). Thus, new approaches are required to complement long-standing techniques for surveying animal biodiversity: analysis of environmental DNA (eDNA) is once such new approach that shows promise for a range of biomonitoring applications (Thomsen & Willerslev, 2015).

Environmental DNA is DNA that is shed from organisms via skin cells, hair, blood, faeces etc. (Ruppert et al., 2019; Taberlet et al., 2012). These small fragments of persistent, but often degraded, single- or multi-source DNA can be collected from the environment, amplified and sequenced, a process known as ‘metabarcoding’, to monitor biodiversity (Ji et al., 2013; Ruppert et al., 2019). All DNA fragments are not preserved equally however, with detectability influenced by a range of factors including rate of DNA shedding, biomass and mobility of the organism the DNA comes from, the type of substrate from which the DNA is extracted, interaction of the organism with the substrate, and a variety of environmental factors such as temperature, rainfall, UV etc. (as reviewed in Newton et al. 2025). Many substrates have been investigated as sources of vertebrate eDNA including bulk environmental samples such as whole insects (Fernandes et al., 2023), faeces (van der Heyde et al. 2021; Nørgaard et al., 2021), flowers (Newton et al., 2023), water (Foote et al., 2012; McDonald et al. 2022), soil (Andersen et al., 2012), vegetation swabs (Lynggaard et al. 2023) and recently, air (Clare et al., 2022; Johnson et al., 2023; Lynggaard et al., 2022). Whilst substrates like water have been extensively studied for vertebrate eDNA detection and are already being employed in marine biodiversity surveys and monitoring (Gilbey et al., 2021; Sigsgaard et al., 2020), analyses of air as a substrate is still in the developmental phase, with current research focused on optimising collection and processing techniques.

Air is increasingly being recognised as a biologically rich source of taxonomically diverse eDNA, including that from vertebrates (Clare et al., 2022; Johnson et al., 2023; Lynggaard et al., 2022). Air contains microscopic particles that have been released into the environment as bioaerosols (Roger et al., 2022) including eDNA which is referred to as ‘airborne eDNA’ or ‘airDNA’ (Clare et al., 2021; Longhi et al., 2009). Until recently, airborne eDNA studies focused on detecting allergens and fungal pathogens from the air, primarily to assess impacts on human health (Mohanty et al., 2017; Kraaijeveld et al., 2015). Multiple proof-of-concept studies that explore air as a medium for the detection of single species and even whole communities of terrestrial species have now been published, including mammals (Clare et al., 2021; Lynggaard et al., 2024), non-anemophilous plants (i.e. plants that do not rely on wind for pollination; Johnson et al., 2019) and insects (Roger et al., 2022). A key benefit of sampling airborne eDNA is that air (unlike other substrates) is ubiquitous. However, the ecology (origin, state, transport, and fate) of airborne eDNA is not yet well understood and is probably affected by both the species targeted and environmental factors (Johnson & Barnes, 2024). Thus, analysis of airborne eDNA remains a nascent method and, like any emerging methodology, requires through investigation to optimise its application.

Multiple approaches have been trialled to effectively capture airborne DNA for vertebrate detections, yet no standardised sampling method has been developed. Active air sampling involves drawing air through a filter membrane using a power source (Bohmann & Lynggaard, 2023; Lynggaard et al. 2024) such as a vacuum or via a fan. However, these samplers can be expensive and/or impractical to run for long periods, particularly in remote areas. Alternatively, passive air sampling relies on the deposition of air particles onto a stationary surface without an externally generated airflow (Bohmann & Lynggaard, 2023). Both passive (Johnson et al., 2023) and active (Clare et al., 2022; Garrett et al., 2023; Lynggaard et al., 2024) air filtering methods have proven effective, though active air sampling has been more widely adopted to date (Johnson & Barnes, 2024). Passive filters are advantageous in that they can be left out for longer periods and are generally less expensive than active ones (Johnson & Barnes, 2024). Additionally, passive filters can be harnessed from novel sources already present in the environment; for example, sampling spiderwebs allowed eDNA detection of vertebrate species in a natural woodland setting and in a zoo (Newton et al., 2024). Adapting existing infrastructure like car cabin filters (Hurley et al., 2019) and air monitoring systems (Littlefair et al., 2023) shows promise for rapid high-throughput sampling of large areas.

Here we evaluated a novel method for airborne DNA sampling by fixing a customised air filter to the roof of a moving car and driving over 250km of transects through woodland and dryland cropping areas in the south-west of Western Australia. We compared the vertebrate biodiversity we detected for these natural and highly modified environments using our car-mounted filters to that obtained from short- and long-term passive filters placed along the same transects. A mobile air sampling method has the potential to more efficiently sample over larger distances compared with other air filtration methods. Similar approaches have been employed for bacteria, plants and vertebrates using a plane (Métris & Métris, 2023) and for marine eukaryotes using a sea vessel (Câmara et al., 2023), but to our knowledge no one has previously made use of a moving land vehicle.

We hypothesised that mobile and long-term passive sample collection would detect higher species richness than would the short-term passive method, and that species richness would be higher in woodland compared to dryland cropping habitats. We expected that the two land uses will comprise distinct communities (i.e. native species in the woodland and domestic species in the dryland cropping areas). Ultimately, this study aimed to expand on the foundational research of airborne eDNA analysis by methodically testing and comparing multiple collection techniques.

## METHODS

### Study Site

This study was conducted in and around the Dryandra Woodland National Park (32.78°S, 116.97°E), 170 km south-east of Perth, Western Australia (Figure 1). Dryandra Woodland is the largest natural vegetation remnant in the Wheatbelt region (∼ 280km^2^) and is characterised by open eucalypt woodland (*Eucalyptus wandoo, E. accedens, E. lane-poolei*), with an understory of sheaoak (*Allocasuarina huegeliana*) and banksia (*Banksia nobilis, B. armata*; Angel & Bradley, 2022). The woodland supports a diverse range of vertebrate species, many of which are of conservation significance (Friend et al. 1995). It is, however, highly fragmented, comprising 17 blocks of remnant vegetation, which are now surrounded by cleared properties primarily used for cereal cultivation, and for sheep farming (Saunders, 1989; Hobbs 1993).

**Figure 1.**
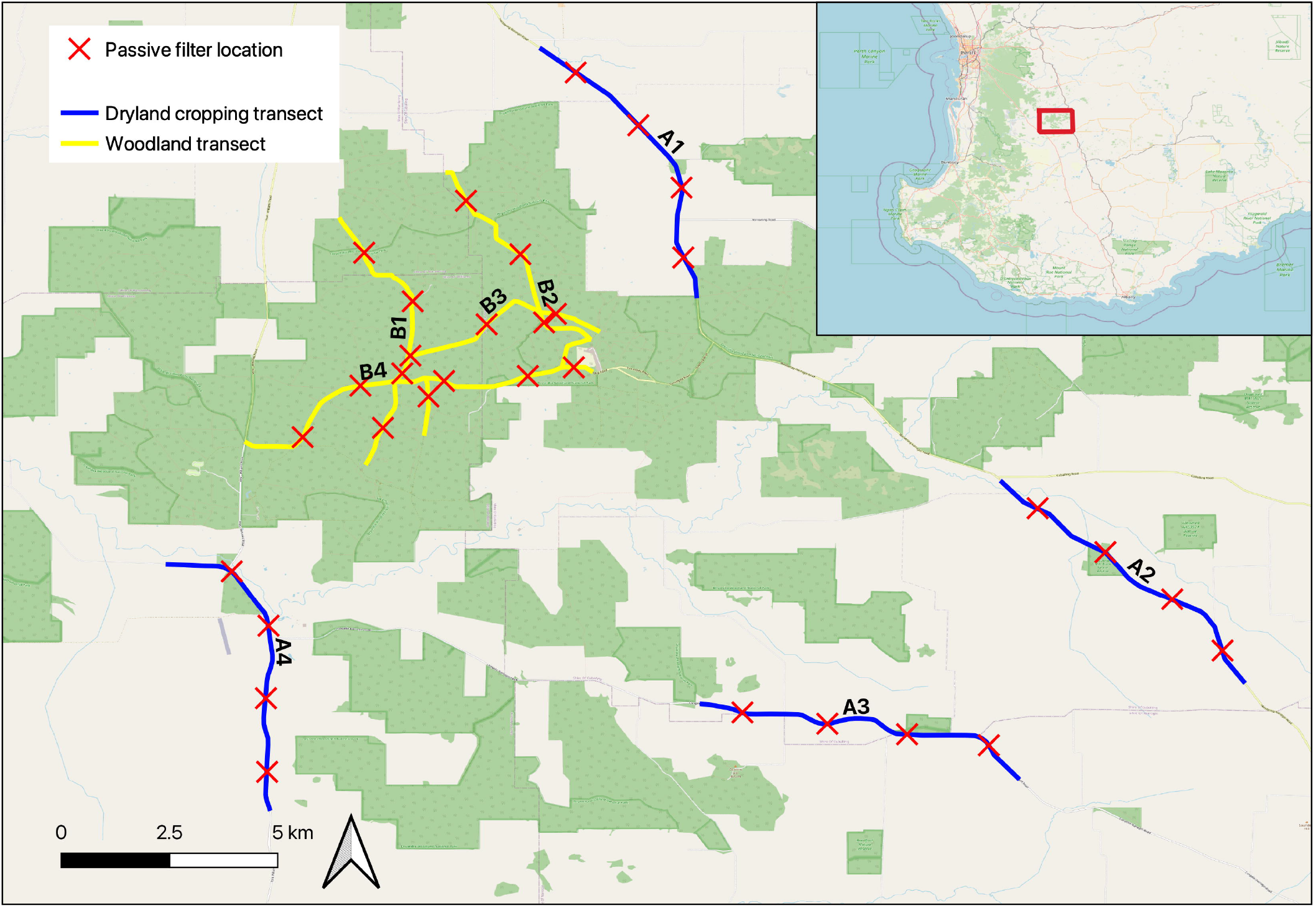
Dryandra Woodland National Park is in the south-west of Western Australia. Airborne DNA transects were each 8 km long; **dryland cropping** transects (A) are shown in blue, woodland transects (B) in yellow. The locations of passive filter placements are marked by red crosses. The same transects were driven on two consecutive days during May 2024

Eight stretches of road spanning 8 km each were selected as ‘transects’ for sampling purposes, four inside the main block of Dryandra Woodland (area 120 km^2^) and four moving from adjacent to the woodland into adjoining dryland cropping areas (Figure 1). The same transects were driven on two days. Due to the limited number of accessible roads within the woodland, these transects overlapped at some points.

### Sample Collection

The vehicle-mounted filter was designed to enclose a strip of filter paper between two pieces of acrylic plastic through which four circles were laser-cut (Figure 2). This ensured that four biological replicates were obtained from each car filter. Prior to sampling, in designated clean laboratories at Curtin University, Perth, Australia, all materials were sterilised in a 10% bleach solution for a minimum of 10 minutes, thoroughly rinsed with deionised water, dried within a laminar flow hood, and UV-sterilized for 15 minutes. The filter media used for air sampling was MERV 13 pleatable synthetic filter paper (Pearl Filtration, Australia). Prior to being sterilised, filter paper was cut to size, either into small 40 mm x 40 mm squares (for passive sampling) or large 50 mm x 300 mm strips (for mobile sampling). Once sterilised, the small square filter papers were attached to sterilised bulldog clips and placed in individual plastic zip-lock bags until sampling. The car-mounted air filters were assembled by screwing two acrylic plates together with the large strip of filter paper in between. These were also placed in individual plastic bags until deployment.

**Figure 2.**
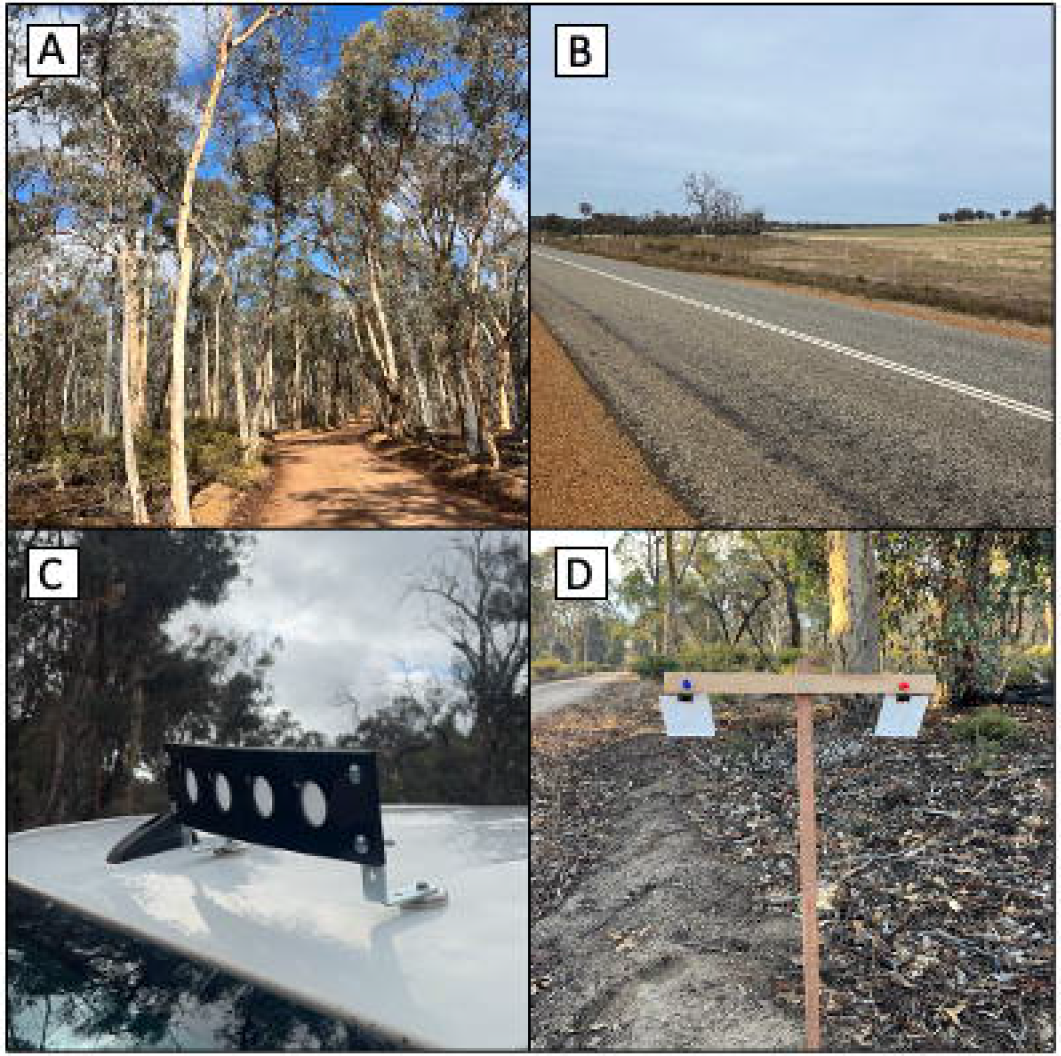
(A) Transect within Dryandra Woodland; (B) Transect on the outskirts of Dryandra woodland, surrounded by dryland cropping; (C) Filter fixed to the roof of a car whilst driving along each transect; (D) Filters placed out along each transect to passively collect air samples. One filter paper at each site was left out for approximately 40 minutes simultaneous to mobile sampling (on each of the first two days) whilst the other was left in place for six days.

Samples were collected between the 11^th^ and 16^th^ of May 2024. Mobile sampling and short-term passive (SP) sampling took place on the first two days (11^th^ and 12^th^ of May; day one and day two respectively), whilst long-term passive (LP) filters were deployed on the 11^th^ of May and collected on the 16^th^ of May. A total of 160 samples were collected during the sampling period which were stratified according to sampling method, land use and sampling day (Table 1).

**Table 1.** Experimental layout of sample collection used in this study. Number of samples are split for mobile and short-term passive into day 1 and day 2 (day 1/day 2).

| Sampling Method | Land use |  | Total Samples |
| --- | --- | --- | --- |
|  | Nature conservation | Dryland Cropping |  |
| Mobile | 16/16 | 16/16 | 64 |
| Short-term passive | 16/16 | 16/16 | 64 |
| Long-term passive | 16 | 16 | 32 |
| **Total Samples** | 80 | 80 | 160 |

At each transect, on both day one and day two, a pre-assembled mobile filter plates was removed from its plastic bag and mounted onto the roof of the car using L-shape brackets and rare-earth magnets (Figure 2). Transects were driven at a speed of ∼30km hour^-1^ and therefore the filter remained on the car for approximately 40 min (the time taken to drive the length of the transect twice i.e. 16 km). After 16 km, mobile filter plates were removed from the car and placed on aluminium foil and each filter (n = 4) was then cut from the mobile filter plates with a new scalpel blade and placed into individual sterile plastic bags. Two separate cars were used to drive the transects in a random order in each land use to minimise the effects of time of day and weather.

Simultaneously, and along the same transects as the mobile sampling, a total of 64 SP samples (32 in each land use across two days; Table 1) were deployed. These consisted of four small filter papers placed evenly along the length of each transect, the first 1 km from the start of each transect and then every 2 kilometres thereafter (Figure 1). The filter papers held in bulldog clips were attached to a wooden stake using push pins (Figure 2). After completing a mobile transect, SP filter papers were collected and placed into sterile plastic bags. The LP filter papers were also deployed along each transect (n = 32 samples; 4 per transect) on the 11th of May, in the same locations as SP samples (Figure 2). These were collected on the 16^th^ of May. All samples were stored on ice after collection, for up to 48 hours, before being transferred to the freezer (-20°C) until DNA extraction.

To minimise contamination of samples, sterile gloves were worn when handling filter papers. Each scalpel blade was used once only and then discarded. Two sterile, unused filters (pre-treated in the same way as all others) were transported to and from the sampling site but remained sealed for the duration of sampling to act as field controls. These were processed along with all other samples, and all samples were processed individually.

### Sample Processing

Sample processing took place in the Trace and Environmental DNA (TrEnD) laboratories at Curtin University, Perth, Western Australia. All air filters were defrosted and transferred from their plastic bags into 2 ml Eppendorf tubes using sterile tweezers. Air filters cut out of the car-mounted mobile filter plates were placed in whole, whilst passive air filters were first cut in half (due to their size) and then placed in the tube. DNA was extracted from air filters using the DNeasy Blood and Tissue Kit (QIAGEN), with the following modifications to standard protocol: either 1000 or 1200 μl of ATL lysis buffer, 60 μl of proteinase K, and overnight digestion at 56°C. Blank extraction controls (n = 6) containing reagents only were processed alongside each batch of extractions to detect any cross-contamination. Resulting eDNA extracts (100 μl) were then stored at - 20°C.

Quantitative polymerase chain reaction (qPCR) was performed on a representative subsample at 3 dilutions (neat, 1/5, 1/10) to quantify DNA present in each sample and assess for inhibition. The 12S-V5 general vertebrate primer (F2: 5′- TAGAACAGGCTCCTCTAG-3′; R2: 5′-TTAGATACCCCACTATGC-3′; Riaz et al., 2011) was used to amplify a ∼98□bp fragment of the 12S rRNA gene. The 12S-V5 primer was selected as it is the most used primer in vertebrate eDNA studies and has been shown to work effectively in eDNA studies in Western Australia (Newton et al., 2024; Ryan et al., 2022; van der Heyde et al., 2020). Extracts were made up in 25 μl reaction volumes using the following reagents: 2.5□mM MgCl_2_, 1□×□PCR Gold Buffer, 0.25□mM of dNTPs, 0.4□mg/mL BSA, 0.4 µM/L of each primer, 0.6 μl of a 1:10000 solution of SYBR Green dye, 1U AmpliTaq Gold, 2□μl of DNA and the remaining volume with distilled water. Extracts were amplified on a StepOnePlus Real-Time PCR System (Applied Biosystems) under the following conditions for 12S-V5: initial denaturation at 95°C for 5□min, followed by 50□cycles of 30□s at 95, 52°C for 30□s and 2□min at 72°C, with a final extension for 10□min at 72°C. Positive controls and non-template controls (NTC) were run with each qPCR batch. Samples did not exhibit PCR inhibition, and therefore it was determined that all samples would be run neat (i.e. without dilution) for fusion tagging.

Quantitative PCR was also performed on a sub-sample both with and without a human blocking primer (5′–3′ TACCCCACTATGCTTAGCCCTAAACCTCAACAGTTAAATC – spacerC3) (Calvignac-Spencer et al., 2013) to determine whether it should be used in the final assay. Human-specific blocking primers aim to reduce the chance of obtaining false negatives (i.e. eDNA templates swamped with human DNA and hence not amplifying) by ‘blocking’ the amplification of contaminant human DNA during PCR (Boessenkool et al., 2012). Samples were run using the same conditions as described above, however resulted in very late amplification or amplification failure in many of the samples. Based off these results, it was decided that human blocking primers would not be used in the fusion-tagged assay.

Each sample was assigned a fusion-tagged primer, consisting of Illumina compatible sequencing adaptors, and a unique (6-8 bp) multiplex identifier tag (MID-tag) combination. Fusion-tagged qPCR reactions were prepared in the TRACE facility at Curtin University using an automated QIAgility robotics platform (QIAGEN). Fusion-tagged qPCR was carried out in duplicate and with the same reagents and cycling conditions described above.

Fusion-tagged amplicons were pooled according to qPCR ΔRn values, in approximately equimolar concentrations (within 5000 ΔRn). Minipools were then screened using a QIAxcel (QIAGEN), resulting in the creation of two libraries (one > 10 ng μL-1, one < 8 ng μL-1). Both libraries were size-selected to 150-300 bp using a Pippin-Prep 2% Agarose Gel Cassette (Sage Science, Beverly, USA) to remove non-target amplicons and primer-dimer. They were then purified using a QIAquick PCR Purification Kit (QIAGEN) and quantified using a Qubit 4.0 Fluorometer (Invitrogen). Sequencing was performed on an Illumina NextSeq platform (Illumina) by loading the library onto a 300 cycle NextSeq reagent cartridge with a P1 Flow Cell for single-end sequencing.

### Bioinformatics

Sequencing data was analysed with the eDNAFlow bioinformatics pipeline of Mousavi-Derazmahalleh et al., (2021) via the Setonix supercomputer at the Pawsey Supercomputing Centre in Kensington, Western Australia. Initial sequence filtering, zero-radius operational taxonomic unit (ZOTU) formation, post-clustering curation and taxonomic assignment were performed during this stage as follows. Sequences were quality-checked using FASTQC (Andrews, 2010) and quality filtered using AdapterRemoval v2 (Phred quality score < 20) (Schubert et al., 2016). Sequences were then demultiplexed using OBITOOLS (Boyer et al., 2016) and those shorter than the minimum length of 90 bp were removed. ZOTUs with a minimum sequence abundance of 4 were formed using the USEARCH Unoise3 algorithm (Edgar, 2016). ZOTUs were then queried against the GenBank (NCBI) nucleotide reference database using BLASTN (Altschul et al., 1990) with 100% query coverage and 95% identity. Post-clustering curation was performed using LULU (--minMatch_lulu ‘97’)(Frøslev et al., 2017). eDNAFlow LCA script (Mousavi-Derazmahalleh et al., 2021) was then used to assign ZOTUs to their lowest common ancestor (LCA); when the percentage identity between two hits differed by ≤ 0.5% with 100% query cover and 98% identity, taxonomic identification was assigned to a ZOTU.

All ZOTUs were then manually verified using open access biodiversity distribution data (Atlas of Living Australia, 2021). When a query sequence matched more than one species (and was therefore dropped to genus or family level), the sequence was manually checked using Geneious Prime 2023.0.3 to determine whether it could be assigned based on distribution data. If the query sequence matched with only one non locally occurring species (i.e. Eastern Rosella) unlikely to reside in the area, then it was either 1) dropped to the lowest taxonomic level that included related species known to inhabit the area (if no related species were present, then it was removed from the dataset), or 2) reassigned to a closely related species within the same genus (i.e. Western Rosella) known to occur in the area. All human ZOTUs and ZOTUs corresponding to fish sequences were also removed from the dataset.

Two methods (Drake et al., 2022) were then used for further data filtering using the ‘phyloseq’ package in RStudio v.4.2.0 (R Core Team, 2020). Firstly, we filtered out all reads counts < 5 reads based on the observation of ZOTUs in the positive and negative controls. Then we removed read counts that made up < 0.01% of the read abundance of each sample.

### Statistics

All values are presented as mean ± standard error, unless stated otherwise. Numbers of taxa detected by the various sampling methodologies were compared with a uniform distribution goodness of fit test, and the proportion of unique species with a two-way contingency table. A Yates correction was applied when the degrees of freedom = 1, using Statisti*XL* V2.0 (www.statistiXL.com).

To determine whether sampling method, land use or sampling day influenced community composition, a permutational multivariate analysis of variance (PERMANOVA) with 999 permutations was conducted on a Jaccard similarity matrix calculated from the presence/absence of species in each sample, using Primer +7 (Clarke & Gorley, 2015). To visualise compositional differences between sampling method, land use and sampling day, the Jaccard similarity resemblance matrix was subjected to the bootstrap averages routine (Clarke & Gorley, 2015) in n-dimensional metric multi-dimensional scaling (nMDS) space. nMDS ordination plots were constructed from the bootstrapped averages (resampling = 150) for each group of samples (e.g. those for a land use, filter type or sampling day). Superimposed on each plot was a point representing the average of the bootstrapped averages and the bias-corrected region in which 95% of the bootstrapped averages were estimated.

Asymptotic regression rarefaction curves representing the order-free accumulation of mean taxa detections calculated from random permutations of all possible orderings of taxa detections were generated for each sampling method. We then used an EcoTest (package “rareNMtests”; Cayuela et al., 2015) to statistically compare rarefaction curves representing each sampling method for each land use type, testing the null hypothesis that samples were drawn from a single assemblage and any differences in their rarefaction curves reflect only sampling effects. A “leave one out” analysis allowed us to explore which sampling approach drove any observed differences between sampling methods, after Cayuela et al. (2015).

### RESULTS

Overall, a total of 27,049,143 metabarcoding reads (raw sequences) were generated using the 12S-V5 assay from 160 samples and 16 controls (field, extraction, PCR positive/negative, and sequencing controls). Human ZOTUs comprised 23,448,105 reads (86.69% of the total reads) and were subsequently removed, along with ZOTUs unable to be assigned to family level. All ZOTUs corresponding to fish (Actinopteri) were also removed (40,347 reads) as probable contaminants, and because our focus was on detecting terrestrial vertebrate species. Following quality filtering, the 12S-V5 assay yielded 3,557,326 reads (mean per sample = 72,598 ± 35,786) from 160 samples, which represented 49 species, including 15 mammals, 33 birds and one amphibian (Table 2).

**Table 2.**
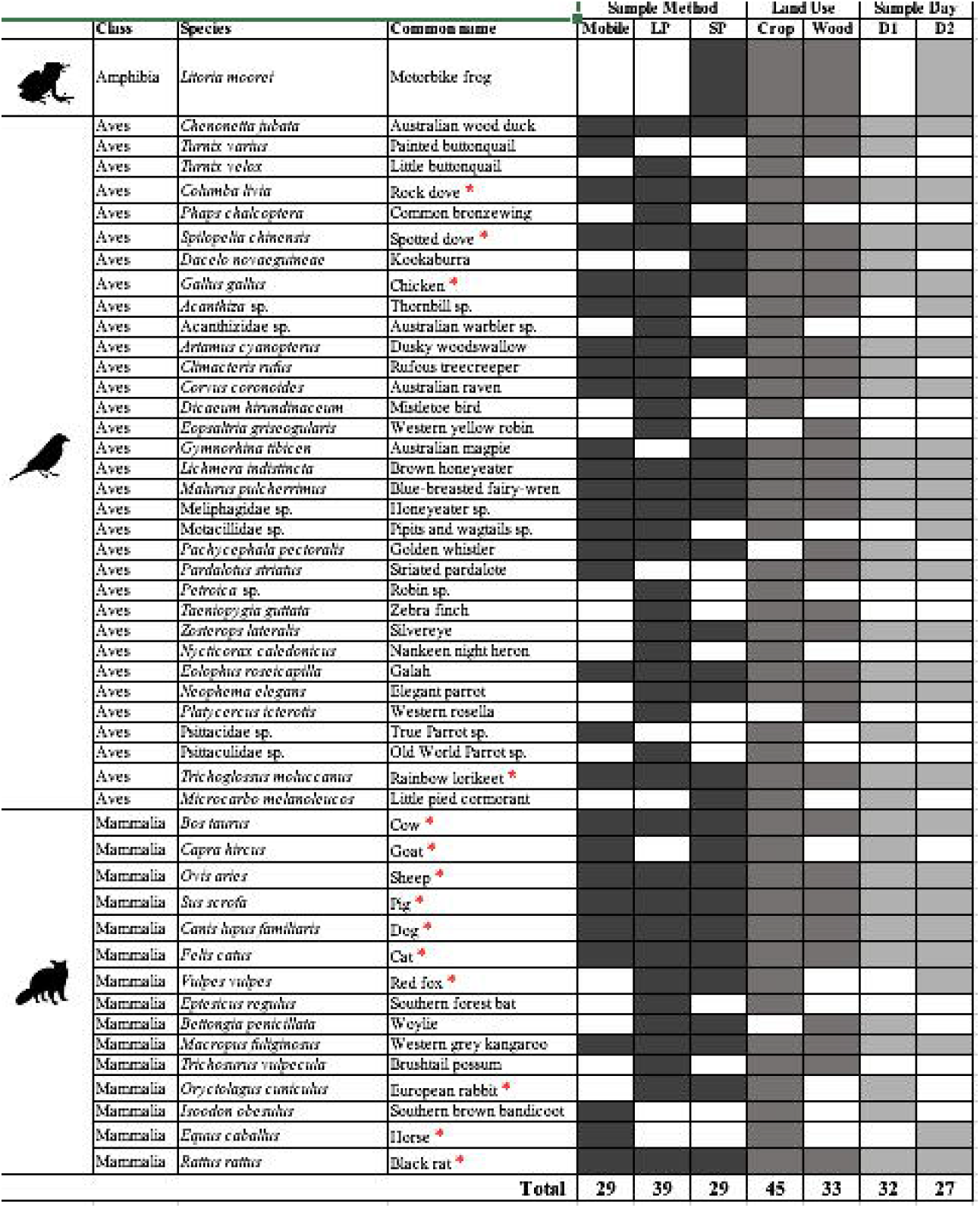
All species detected using the 12S-V5 primer set. Sample methods include car-mounted filters (mobile), long-term passive (LP) and short-term passive (SP) filters. Land uses surrounding the sampling transects were dryland cropping (Crop) or woodland (Wood). Sample day was either day 1 (D1) or day 2 (D2). The ‘Sample Day’ category includes only taxa detected from samples collected using the mobile or SP method. A red star indicates domestic or invasive species (i.e. not native to the area) while a green star indicates the species is found in the south-west of Western Australia, but not specifically known to inhabit Dryandra Woodland. All other species are known inhabitants of Dryandra Woodland.

Thirty-four native or naturalised species known to occur within Dryandra Woodland were detected, with the remaining species consisting of either agricultural/domestic/pest species (n = 12), or other species (i.e. birds) that are known to traverse the south-west region (n = 3). Notable native species detections include the critically endangered woylie (*Bettongia penicillata*), and the conservation priority four (rare, near threatened) Quenda (*Isoodon fusciventer*; Table 2). Despite Dryandra Woodland being well-known for its numbat (*Myrmecobius fasciatus*) population, numbat eDNA was not detected in any samples using the 12S-V5 primer set. Short-beaked echidnas (*Tachyglossus aculeatus*) are also common within Dryandra, but were not detected. The red fox (*Vulpes vulpes*), an introduced predator within the area, and one of the main threats to small native marsupials in Dryandra, was only detected within the dryland cropping area (n = 2 samples), while the woylie was detected only in the woodland (n = 2 samples). Cat (*Felis catus*), another introduced predator, was detected in almost half of the collected samples in both the woodland and dryland cropping transects.

Long term passive sampling yielded 39 taxa, of which 12 were unique detections, which did not differ from the total number of taxa detected by mobile (total taxa = 29, unique detections = 5) and SP (total taxa = 29, unique detections = 3) methods (χ^2^_2_ = 2.1, P = 0.357; Figure 3). The three methodologies detected similar proportions of unique taxa (χ^2^_2_ = 3.0, P = 0.226). Sampling along the agricultural dryland cropping transects detected a total of 45 taxa (16 unique detections) compared to 33 taxa detected in the woodland (with 4 unique detections; Figure 3); land use had no significant effect of numbers of taxa detected (χ^2^_1_ = 1.5, P = 0.213) and there was no difference in the proportion of unique taxa detected (χ^2^ _1_= 2.5, P = 0.115; Figure 3). Of the taxa unique to the agricultural dryland cropping transects, 12 were native animals and four were domestic/introduced species. The four taxa identified only in the woodland are all native to the south-west of WA (Table 2). Sampling day had no effect on the number of taxa detected (χ^2^ _1_= 0.27, P = 0.603) with 32 taxa detected on day one and 27 on day two. Community composition was not significantly impacted by sampling method (F_2,154_ = 1.21, *p* = 0.234) or sampling day (F_1_ = 0.314, *p* = 0.987) but differed significantly between land uses (F_1_ = 2.48, *p* = 0.013). Visualisation by nMDS ordination shows no overlap of bootstrapped 95% confidence ellipsoids of the two land uses (Figure 4).

**Figure 3.**
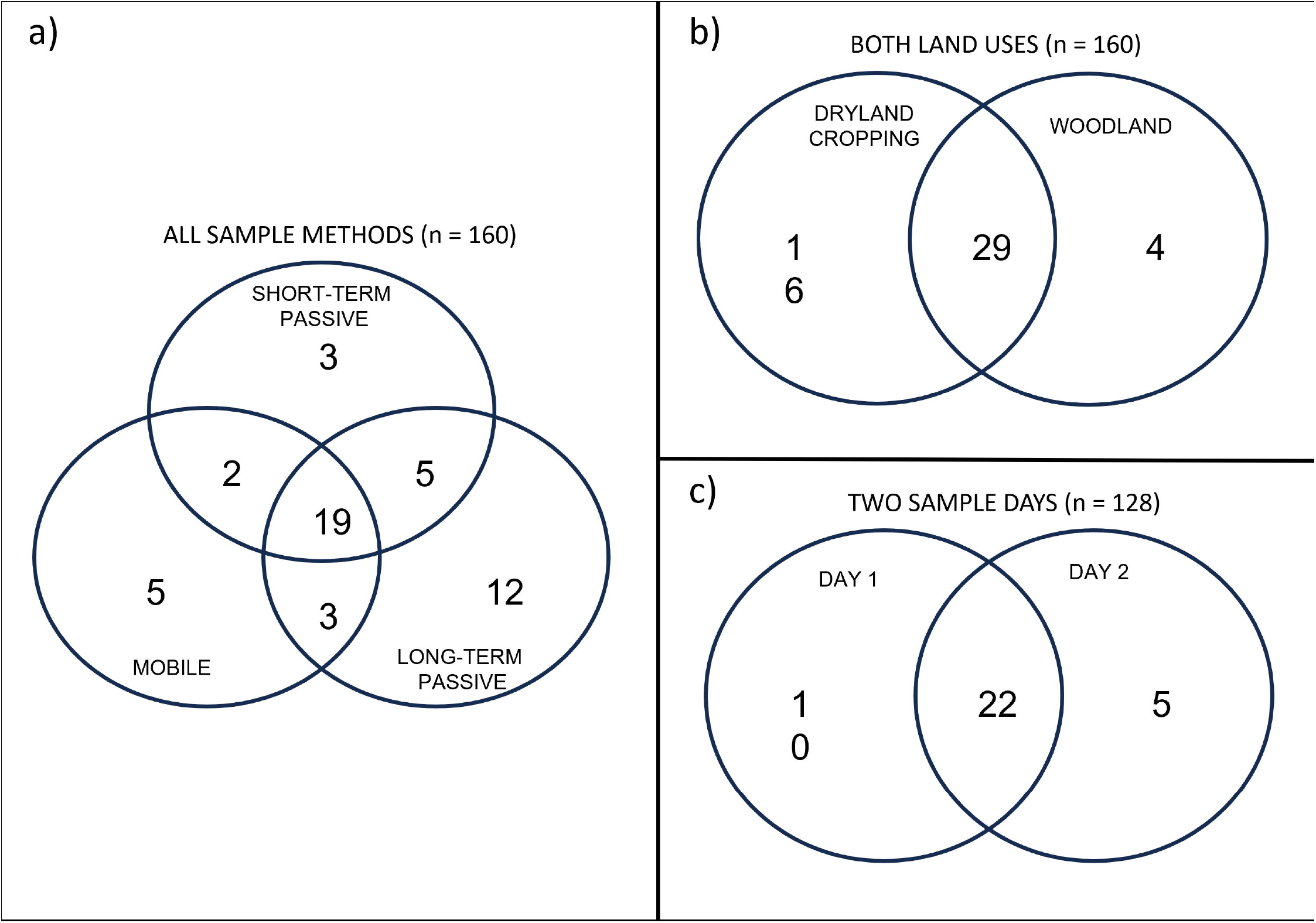
Venn diagrams showing the relationship between species detections using (A) short-term passive (n = 64), long-term passive (n = 32) and mobile (n = 64) sampling methods, (B) dryland cropping (n = 80) and woodland (n = 80) and (C) sample day 1 (n = 64) and sample day 2 (n = 64).

**Figure 4.**
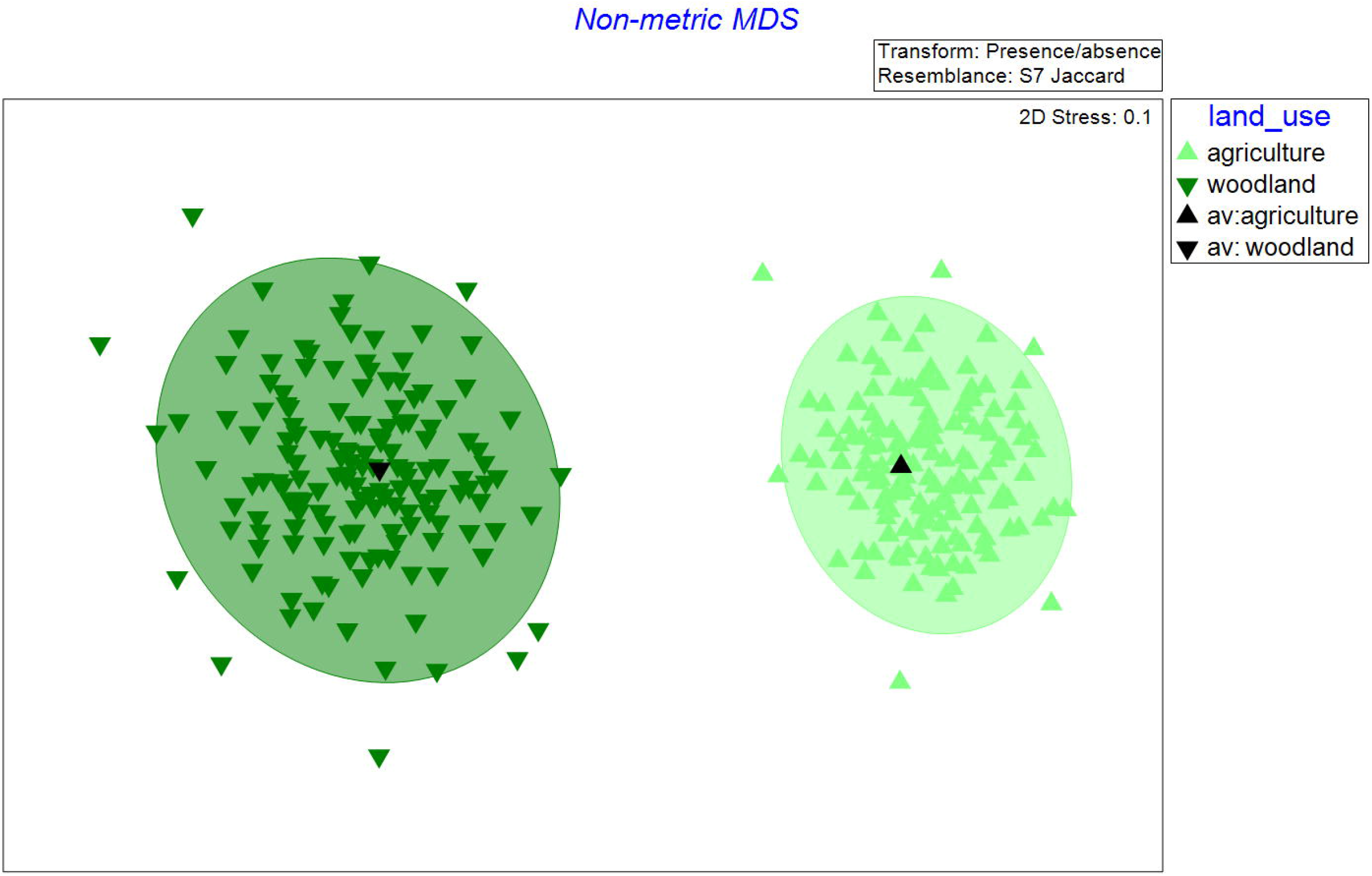
nMDS plot showing community composition for woodland (dark green, n = 80) and dryland cropping (light green, n = 80) land uses. Plot visualises a Jaccard similarity matrix calculated from the presence/absence of species in each sample. Black triangles represent the average bootstrapped value, and the ellipsoid is the 95% confidence interval for each factor.

Rarefaction EcoTests indicate overall differences for species accumulation curves obtained using the different sampling methods (z□=□104; p□=□0.005; Figure 5), but no differences between dryland cropping and woodland sites (z = 80, p = 0.285). Long and short passive sampling curves differed (z = 264, p = 0.025) but car and short-term passive cures did not (z = 50, p = 0.115). None of the curves approached an asymptote during the sampling duration.

**Figure 5.**
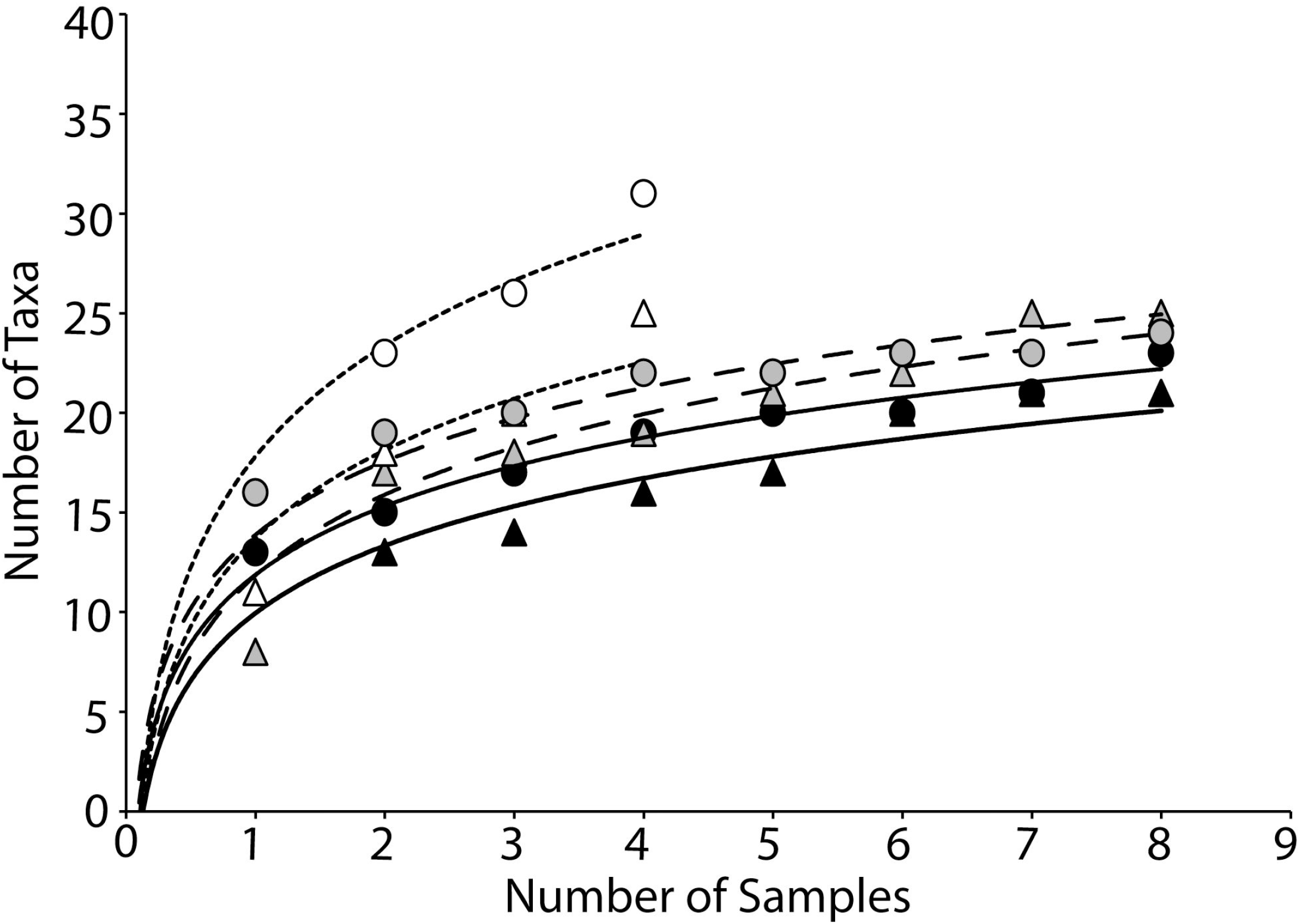
Accumulation curves for vertebrate detections in dryland cropping (circles) and conservation reserve (triangles) areas at Dryandra National Park, Western Australia using mobile (black, solid lines), short-term stationary (grey, long dashed lines) and long-term stationary (white, short, dashed lines) data sampling.

## DISCUSSION

The application of airborne eDNA is quickly gaining traction as a biomonitoring tool (Clare et al., 2022; Johnson et al., 2023; Lynggaard et al., 2022; Tulloch et al., 2025), but only a small number of studies have directly examined it as a source of vertebrate eDNA (Johnson & Barnes, 2024). Here we tested a new method of sampling airborne eDNA in both a natural and an agricultural setting. Our mobile sampling method detected 29 vertebrate species, with a mean species richness per sample comparable to both long- and short-term fixed collection methods. Our mobile sampling method detected five unique species (striated pardalote, a parrot sp., painted buttonquail, horse, and quenda) not detected by the other sampling approaches, including one mammal (the quenda) of conservation concern. Our mobile airborne DNA collection method shows promise for enhancing biodiversity monitoring within terrestrial ecosystems by providing an efficient way to sample large areas.

Due to the relatively recent emergence of air as a substrate for terrestrial vertebrate eDNA sampling, there are no established ‘best practices’ for collection of airborne eDNA. However, similar to our findings for vertebrate taxa, multiple studies focusing on macrobial airborne eDNA have reported that collection method does not impact the number of taxa detected (Garrett et al., 2023; Lynggaard et al., 2024). With the sampling methods employed here, our results suggest that it is the exposure time and the number of samples, rather than the sampling method itself, that determines the accumulation of species recorded in an area.

Interestingly, biological replicates collected on the same mobile air filter (n = 4 per mobile filter) did not consistently detect the same species, highlighting potential variability in airborne DNA distribution at very fine spatial scales (e.g. Harnpicharnchai et al. 2023). As pointed out by Claire et al. (2021), airborne DNA analyses suggest that DNA in the air “will show spatial–temporal confinement just as eDNA in water has been found to be spatially confined”. The detection of uniquely identified species on each sampling day (day 1 = 10 unique taxa; day 2 = 5 unique taxa), even from biological replicates exposed simultaneously on a single mobile sampling transect, suggests that large filter surfaces and/or spatial and temporal replication is necessary to capture a reasonable representation of species present in a given area. Consistent with findings from other eDNA studies (Koziol et al., 2019; Ryan et al., 2022; van der Heyde et al., 2020), and indeed other monitoring approaches in general (e.g. Thompson and Thompson 2007; Thompson et al. 2003, 2007), our results reinforce the positive effects of increased sampling effort, both in terms of sample size (i.e. placing more filters on the car, leaving samples in place longer for passive samples, or traversing transects multiple times for mobile samples) and potentially also number of sampled substrates, to detect a greater proportion of the taxa present.

Our three methods for airborne eDNA sampling, employed for one week, allowed us to detect approximately 20% of the vertebrate species known to be present in the area. However, considering the 49 taxa detected included domestic animals, more than three-quarters of the species known to inhabit Dryandra Woodland remained undetected by any of the sampling methods. While it is unlikely that a single survey/monitoring method will ever detect all species within an area, several factors may account for this low detection rate. Notably, we failed to detect any dasyurmorph marsupials e.g. chuditch, yellow-footed antechinus, red-tailed phascogale or numbat, nor the monotreme echidna. This may be due to biases in the substrate sampled; arboreal species such as the phascogale and antechinus may be more easily detected in tree swabs (Newton et al., 2022), while digging species like the echidna may be detectable by sampling disturbed soil. Another factor that can influence species detection is the significant gaps that remain in the reference sequence database (Keck et al., 2023). Of the 201 native vertebrate species recorded in the Dryandra (Atlas of Living Australia, 2021), 79 do not have 12S DNA sequences on GenBank. For example, 12S DNA sequences from the western rosella (*Platycercus icterotis*), a moderately common bird in the south-west of Western Australia, does not appear within the GenBank database (Benson et al., 2009) – one of the oldest and most used open-source DNA sequence databases (Keck et al., 2023). This is also true of many other species that inhabit the south-west region. A final explanation of why the number of species detected by airborne DNA surveys is not the same as the number of species known to occupy the area is that some of the species in the woodland are migratory e.g. many of the Meliphagidae (Recher et al. 2016; Newland & Wooller 1985) and will consequently not be there all the time.

Our approach to analysing airborne DNA could detect significant differences in community composition between woodland and adjacent dryland cropping areas. This suggests that although airborne DNA may potentially disperse further from the source than DNA obtained from more contained substrates such as sediment, hollows or discrete waterbodies (Lynggaard et al., 2024), species detections were not ubiquitous across the landscape, with only 59% of all species detected in both land use types. Seventeen additional taxa were identified in the dryland cropping zone compared to the woodland, including both introduced/domestic e.g. horses (*Equus caballus*), rabbits (*Oryctolagus cuniculus)*, and foxes, and native e.g. southern forest bat (*Vespadelus regulus*), quenda, and various native bird species. The highly fragmented nature of Dryandra Woodland means that native species reliant on remnant native vegetation often move through the dryland cropping matrix surrounding the remnants (Bennett, 1991), and patches of woodland both in paddocks and on the roadsides were common along the dryland cropping transects (Figure 1). It should also be noted that 11 of the 16 species only detected in the dryland cropping area were volant; presumably flight facilitates movement throughout the landscape, including highly modified land use types (Withers et al. 2004; Doherty and Driscoll 2018). This issue could be further examined by conducting additional studies with dryland cropping transects located farther from the woodland.

It is well established that substrates perform differentially, and some are more conducive to detecting certain taxa (van der Heyde et al., 2020). Like previous airborne DNA studies (Lynggaard et al., 2024; Polling et al., 2024), we found birds to be the most frequently detected class (33 out of 49 taxa). While an effective substrate for detecting birds and mammals, air appears to be considerably less suitable for detecting amphibians and reptiles (Garrett et al., 2023; Lynggaard et al., 2024; Lynggaard et al., 2022). There are 26 reptiles and 4 amphibians known to inhabit Dryandra Woodland but we didn’t detect any reptiles and only one amphibian. This is not surprising as amphibians hide most of the time and some live in water, and reptiles are believed to shed very little eDNA (Nordstrom et al., 2022). Improved reference databases, PCR primer optimization, and sampling a combination of substrate types is likely required to capture the full range of vertebrate taxa present.

Contamination is a significant issue for airborne DNA collection (Johnson & Barnes, 2024), primarily from humans during deployment, collection and lab analysis. We assigned 86.6% of prefiltered reads to human ZOTUs, much higher than in non-airborne DNA studies, which typically show values between 14% -62% in water (Kelly et al. 2014; MacDonald et al. 2023). However, similar high human DNA values have been reported for other airborne DNA studies (Lynggaard et al., 2024; Clare et al., 2021). Human blocking primers can prevent this issue (Boessenkool et al., 2012), but during the early phase of our research, they reduced or prevented amplification in many samples (Figures S1 and S2) and were not used in final analyses. Fortunately, although human DNA dominated total reads, at-depth sequencing on the Illumina NextSeq platform without blocking primers, still yielded over 3 million non-human reads post-filtering, from a raw dataset of ∼26 million reads.

All three methods we used for sampling airborne DNA successfully detected vertebrate taxa. We have, therefore, demonstrated that sampling using a moving vehicle is feasible and can be as effective as other passive airborne DNA sampling methods. Therefore, this study further validates the viability of airborne eDNA for detecting vertebrate species in biomonitoring studies and demonstrates its potential application to other mobile sampling approaches, such as drone-based sampling. From our observations we recommend that mobile sampling should take place over multiple days, use multiple transects, and collect multiple samples per transect. Additionally, stringent contamination controls (gloves, mask etc.) should be employed to limit the influence of human DNA on vertebrate detections. We conclude that the mobile air sampler presents a practical and effective tool for airborne eDNA collection, particularly in scenarios where sampling time is limited and when detecting all species at a location is not essential, and may be of particular benefit for rapidly sampling large areas to obtain a “snapshot” of biodiversity. Further studies are required to examine the effectiveness of car-mounted airborne DNA sampling for other taxa like invertebrates, plants and fungi.

